# Sex-specific GABAergic microcircuits that switch vulnerability into resilience to stress and reverse the effects of chronic stress exposure

**DOI:** 10.1101/2024.07.09.602716

**Authors:** Tong Jiang, Mengyang Feng, Alexander Hutsell, Bernhard Luscher

**Affiliations:** Department of Biology, Pennsylvania State University, University Park, PA 16802.; Department of Biochemistry & Molecular Biology, Pennsylvania State University, University Park, PA 16802; Center for Molecular Investigation of Neurological Disorders (CMIND), The Huck Institutes of the Life Sciences, Pennsylvania State University, University Park, PA 16802; Picower Institute of Learning and Memory, Massachusetts Institute of Technology, 43 Vassar St, Cambridge MA 02139

**Author notes:** **Address for correspondence:** Bernhard Luscher, Ph.D, Department of Biology, Penn State University, 301 Life Sciences Building, University Park, PA 16802.

## Abstract

Clinical and preclinical studies have identified somatostatin (SST)-positive interneurons as key elements that regulate the vulnerability to stress-related psychiatric disorders. Conversely, disinhibition of SST neurons in mice results in resilience to the behavioral effects of chronic stress. Here we established a low-dose chronic chemogenetic protocol to map these changes in positively and negatively motivated behaviors to specific brain regions. AAV-hM3Dq mediated chronic activation of SST neurons in the prelimbic cortex (PLC) had antidepressant drug-like effects on anxiety- and anhedonia-like motivated behaviors in male but not female mice. Analogous manipulation of the ventral hippocampus (vHPC) had such effects in female but not male mice. Moreover, activation of SST neurons in the PLC of male and the vHPC of female mice resulted in stress resilience. Activation of SST neurons in the PLC reversed prior chronic stress-induced defects in motivated behavior in males but was ineffective in females. Conversely, activation of SST neurons in the vHPC reversed chronic stress-induced behavioral alterations in females but not males. Quantitation of c-Fos^+^ and FosB^+^ neurons in chronic stress-exposed mice revealed that chronic activation of SST neurons leads to a paradoxical increase in pyramidal cell activity. Collectively, these data demonstrate that GABAergic microcircuits driven by dendrite targeting interneurons enable sex- and brain-region-specific neural plasticity that promotes stress resilience and reverses stress-induced anxiety- and anhedonia-like motivated behavior. Our studies provide a mechanistic rationale for antidepressant efficacy of dendrite-targeting, low-potency GABA_A_ receptor agonists, independent of sex and despite striking sex differences in the relevant brain substrates.

## INTRODUCTION

Chronic and excessive amounts of stress are key vulnerability factors for virtually all psychiatric disorders, including especially Major Depressive Disorder (MDD) and Post-traumatic Stress Disorder (PTSD). These disorders are characterized by changes in negatively and positively regulated motivated behavior that manifest as heightened anxiety and anhedonia, respectively. Comparable changes in motivated behavior are observed in mice exposed to chronic stress. At the molecular level, MDD and chronic stress are associated with diverse deficits in GABAergic inhibition ^1, 2^. Moreover, molecular genetic studies in mice indicate that reduced GABAergic inhibitory function results in anxiety- and anhedonia-like changes in negatively and positively regulated motivated behaviors ^3–5^. By extension, these studies in patients and mice suggest that reduced GABAergic inhibition contributes causally to stress-associated psychiatric disorders. Conversely, successful antidepressant drug treatments of patients and mice result in normalization of GABA concentrations and restoration of impaired synaptic transmission ^6–11^ along with reversal of the detrimental behavioral effects of chronic stress exposure ^10, 12^. Together these studies predict that GABA itself has antidepressant drug-like properties that promote stress resilience ^13^. A separate body of work suggests that chronic stress and MDD are associated with reduced activity of frontal cortical output neurons ^14^ and reduced functional connectivity ^15–17^, while selective stimulation of pyramidal cells in the medial prefrontal cortex (mPFC) of mice reverses stress-induced defects in motivated behavior ^18–21^. A key question arising from these observations is whether GABAergic inhibition increases or reduces the activity of cortical output cells. Lastly, stress-sensitive psychiatric conditions in general and depressive disorders, in particular, show prominent sex differences in prevalence and disease-associated transcriptomes and suggest that sex differences are involved in the underlying mechanisms ^22, 23^.

In testing whether selective potentiation of GABAergic inhibition at dendrites of pyramidal neurons (PNs) promotes stress resilience, we recently showed that genetically induced global disinhibition and corresponding hyperexcitability of somatostatin (SST)-positive GABAergic interneurons in mice results in behavioral changes that mimic those of antidepressant drug treatment, as well as resilience to chronic stress-induced anxiety^24, 25^. Here we tested whether the emotional behavioral changes of SST neuron-mediated inhibition can be reproduced by chronic DREADD (Designer Receptors Exclusively Activated by Designer Drug)-mediated activation of SST neurons and mapped to specific brain regions in adulthood, whether any such effects are sex-specific, and whether chronic stress-induced alterations in motivated behavior can be reversed by chronic activation of SST^+^ interneurons in specific brain regions. We focused on the prelimbic cortex (PLC) and ventral hippocampus (vHPC) as brain regions that are highly stress-sensitive and important for the top-down control of positively and negatively regulated motivated behavior ^26^ ^27^. In addition, we asked how activation of SST^+^ neurons associated with anxiolytic and anti-anhedonic behavioral consequences affects the activity of their putative pyramidal target cells in vivo.

## MATERIALS AND METHODS

### Animals

SSTCre (also known as *Sst^tm2.1(cre)Zjh^*/J, JAX stock #013044), PVCre (B6.Cg-*Pvalb^tm1.1(cre)Aibs^*/J, Stock No: 012358) mice and C57BL/6J mice (BL6, Stock No. 000664) were obtained from Jackson Laboratory (Bar Harbor, MN). SSTCre and PVCre breeder mice were maintained as heterozygotes and routinely backcrossed with wildtype 129X1/SvJ mice (RCC Ltd Biotechnology and Animal Breeding, Füllinsdorf, Switzerland) and BL6 mice, respectively. The breeding colony was kept under a 12:12h light-dark cycle, on corn cob bedding and with food and water ad libitum. Offspring were genotyped by PCR of tail biopsies at the time of weaning and then sorted into gang cages of 5–6 mice per cage, under an inverted 12:12h light-dark cycle. All animal experiments were approved by the Institutional Animal Care and Use Committees (IACUC) of the Pennsylvania State University and conducted per guidelines of the National Institutes of Health (NIH).

### Chemogenetic Manipulation

Stereotaxic surgeries of mice were conducted at 8–9 weeks of age. Cre transgenic mice were anesthetized with vaporized isoflurane/O_2_ (5% initiation, 2% maintenance) and mounted on a stereotaxic frame (Stoelting, Wood Dale, IL). The surgical site was shaved and prepared with betadine, scrubbing with ethanol, and an injection of bupivacaine (4 mg/kg, s.c.). A scalp incision was made along the midline of the skull, the skin was pulled to the sides and bilateral drill holes were made, targeting the prelimbic cortex (PLC, AP: 1.8 mm, ML: ± 0.4 mm, DV: 2.2– 2.3 mm) or ventral hippocampus (vHPC, AP: −2.9 mm, ML: ± 2.9 mm, DV: 3.25 mm)^28^. Equal groups of mice were injected bilaterally with Cre-dependent AAV vectors encoding a double-floxed Designer Receptor Exclusively Activated by Designer Drug (DREADD) construct fused to mCherry (pAAV-hSyn-DIO-hM3Dq(Gq)-mCherry, abbreviated here as hM3Dq, #0474-AAV5) or the double floxed control vector carrying a cDNA for mCherry alone (pAAV5-hSyn-DIO-mCherry, abbreviated here as Ctrl or control vector, #50459-AAV5, Addgene, Watertown, MA), using 0.3 μL of virus per site at a titer of > 1–2 x 10¹^3^ vg/mL. The wound was closed with an absorbable polyglycolic acid suture (5/0, CP Medical 421A). The mice were allowed to recover for at least two weeks before the continuation of experiments. The injection sites were examined at the end of experiments using immunofluorescent staining of floating sections for mCherry followed by confocal microscopy or scanning with an Odyssey® CLx imager (LI-COR, Lincoln, NE).

### Stress Treatment

The mice were exposed to chronic variable stress (CVS) as described ^29, 30^, starting on day one with a tail suspension stressor, followed on day two with a restraint stressor, and on day three with exposure to foot shocks, which were then repeated for a total of 15 or 21 days. For tail suspension, the mice’s tails were taped to a metal crossbar placed 35 cm above the bench, for 1 h. For the restraint stressor, the mice were placed into a perforated 50 ml Falcon® tube for 1 h. Foot shock treatment involved exposure to 100 randomly distributed foot shocks (0.45 mA x 3 s) delivered within 1 h, using a maximum of 10 mice per chamber of a two-compartment shuttle box (SanDiego Instruments, San Diego, CA) with the connecting gate closed. After each stressor, the mice were returned to their home cage.

### Treatment with Clozapine-N-Oxide (CNO)

CNO (NIMH Chemical Synthesis and Drug Supply Program) was dissolved at 100 mg/ml in dimethyl sulfoxide and diluted 1:1 with 0.9% saline and loaded into Alzet® mini pumps (Model 1004, DURECT Corp., Cupertino, CA) designed to release drug at a rate of 0.11 μl/h for up to 4 weeks. For a 25 g mouse, the resultant dose of CNO was calculated as 0.22 mg/kg/h. For minipump implantation, the animal’s back below the neck was shaved followed by anesthesia using isoflurane inhalation and a dose of bupivacaine (4 mg/kg, s.c.) at the surgical site. A skin incision was made, and the minipump was inserted into the surgical opening with the cap facing inwards. Wound clips (SAI Infusion Technologies) used to close the wound were removed seven days post-surgery. Minipumps were removed at the time indicated in the Figures. For experiments that involved the removal of minipumps before behavioral testing, the mice were anesthetized as above, and an incision was made close to but not overlapping with the initial incision. The minipumps were then removed using sterile forceps and the wounds were closed using wound clips.

### Behavioral Testing

The mice were singly housed under an inverted 12:12h light-dark cycle, starting one day before the start of behavioral testing with the two sexes kept in separate rooms. Behavioral testing followed the time course indicated in the Figures, using one test per day. Testing was initiated ∼3 h after the lights went off and conducted under red light except for the open field test (**OFT**). The Elevated Plus Maze (**EPM)** apparatus ^24^ consisted of an elevated (40 cm) crossbar with two open and two closed arms (30 x 5 cm). The closed arms were surrounded by 20 cm walls of clear Plexiglas and the edges of the open arms were raised by 2 mm to reduce accidental falling. At the start of the test, the mice were placed into the center square of the maze, facing a closed arm. The behavior was video recorded for 5 min and the numbers of open and closed arm entries and the open arm duration were automatically recorded using EthoVision XT13 (Noldus Information Technologies, Leesburg, VA). The **OFT** ^24^ was used to assess locomotion and behavioral inhibition in a novel arena exposed to bright light (75 lux). The OFT arena was divided evenly into a 5 x 5 grid with the central 3 x 3 grid designated as the center arena. The mice were allowed to freely explore an opaque Plexiglas arena (50 x 50 x 20 cm) for 10 min. The behavior was video recorded and duration and frequency into the center were computed by EthoVision. For the Forced Swim Test (**FST)** ^24^ the mice were placed into the 4 L beaker containing 3 L of 25 ±1°C water and their activity was video recorded for 6 min. The frequency of inactivity, total duration spent inactive, and latency to the first inactivity during the last 5 min of the test were automatically recorded using EthoVision. Bouts of inactivity were set as an activity level ≤ 1.15% of the maximum and lasting ≥ 2 s. A z-scored mobility index that combined the three measures was computed as described ^31^. For the Novelty Suppressed Feeding Test (**NSFT)** ^24^, the mice were acclimated to an empty Petri dish in their home cage while being food-deprived for 18 h before testing. On test day they were transferred to a corner of a novel test arena (50 x 40 x 12 cm) containing wood chip bedding. The latency to start feeding on a food pellet placed on a piece of cotton nesting square (6 x 6 x 0.5 cm) in the center of the arena was hand-scored for maximally 6 min even if no feeding had occurred. The mice were immediately returned to their home cage containing a dry food pellet placed on a Petri dish. The amount of food consumed during 30 min was recorded. The Female Urine Sniffing Test (**FUST)** ^32^ was used to assess changes in positively motivated, rewarding behavior in male mice. The mice were accustomed to a dry sterile cotton-tip applicator (Patterson Veterinary Supply, Saint Paul, MN) in their home cage for 30 min and then exposed to a new water-soaked cotton tip for 3 min. After 45 minutes they were exposed to a fresh female-urine-soaked cotton-tip. The behavior was video-recorded, and the duration spent sniffing the cotton tip during the first 3 min was scored offline. The Sucrose Splash Test (**SSPT)** was used to assess positively motivated, rewarding behavior in female mice ^33, 34^. The mice were transferred briefly to an empty cage and the fur coat of their back was sprayed with 1 mL of 20% sucrose solution using a spray bottle, to stimulate grooming behavior. They were immediately returned to their home cage and the grooming duration was recorded during the first 5 min using a stopwatch.

### Immunohistochemistry

To standardize the arousal state of animals at the time of perfusion, each mouse was transferred to a novel shoebox cage for 10 min, followed exactly 80 min later by deep anesthesia with avertin (250 mg/kg) and transcardial perfusion with 4% paraformaldehyde in phosphate-buffered saline (PBS). The brains were postfixed overnight, and coronal sections (40 µm) were stored in PBS containing 0.02% NaN_3_ at 4°C until analyses. Immunostaining was performed in PBS supplemented with normal goat serum (2.5%), normal donkey serum (2.5%), 0.3% Triton X-100, and primary antibodies (1 h) followed by washes and incubation with appropriate secondary antibodies overnight. To monitor viral expression, sections were stained with rat anti-red fluorescent protein (RFP) (1:800, mAB 5F8, Chromotek) and then IRDye^®^ 800CW goat anti-rat secondary antibody (1:500, LI-COR). Sections mounted onto glass slides were imaged using an Odyssey® CLx scanner (LI-COR). For analyses of the density c-Fos and FosB expressing neurons, the sections were immunostained with rat anti-RFP (1:800, Chromotek, recognizing mCherry) together with guinea pig anti-c-Fos (AB_2619946, 1:3000, SySy), monoclonal rabbit anti-FosB (1:2000, AB_184938, ABCAM) or rabbit anti-parvalbumin (AB_298032, 1:500, SySy), followed by appropriate mixtures of secondary antibodies. The injection sites were imaged with a ZEISS® LSM 800 (20x objective). Z-stacks of 6-14 regions of interest (ROI, 300 x 300 pixels, 10 μm depth) were captured per animal. The densities of c-Fos^+^ and FosB^+^ cells were automatically quantified using ImageJ. The densities of mCherry^+^, PV^+,^ and double positive cells were quantified manually. The densities of c-Fos^+^ and FosB^+^ mCherry^+^ cells were normalized to the densities of mCherry^+^ cells in the respective image stacks to adjust for variability in the infection density.

### Statistics

Statistical testing was performed using Prism 10 software (GraphPad, La Jolla, CA). Simple two-sample comparisons were done by t-test or, if the data failed the normality test (D’Agostino & Pearson), by Mann-Whitney test. Multiple comparisons were done by 1-way or 2-way ANOVA followed by post hoc Tukey or Sidak test, respectively as indicated in Figure legends. Data from mistargeted mice and statistical outliers (ROUT method, Q = 1%) were removed.

## RESULTS

### Chronic activation of SST neurons of the prelimbic cortex has anxiolytic- and antidepressant-like effects on motivated behavior in male but not female mice

We previously showed that global disinhibition of SST neurons in SSTCre:γ2^f/f^ mice results in behavior that mimics the effects of antidepressant drug treatment in both male and female mice. ^24^. SST neurons of SSTCre:γ2^f/f^ mice are hyperexcitable due to loss of GABAergic inhibitory synaptic input and a significant increase in input resistance ^24^. Moreover, SST neurons are present throughout the brain as well as in diverse peripheral tissues, and the mutation is induced during embryonic development. This raised the question of whether the behavioral phenotype of SSTCre:γ2^f/f^ mice can be reproduced by increasing the excitability of SST neurons in specific brain regions in adulthood. To address this question, we injected the prelimbic cortex (PLC) of male SSTCre mice (8-10 weeks of age) with a Cre-dependent AAV-hM3Dq-mCherry vector (hM3Dq) or a Cre-dependent AAV-mCherry control vector (Ctrl) and, after allowing three weeks for virus expression, implanted all animals with clozapine-N-oxide (CNO)-releasing osmotic minipumps, regardless of virus injected. We aimed to mimic the hyperexcitability of SST neurons observed in SSTCre:γ2^f/f^ mice in AAV-hM3Dq-manipulated SSTCre mice with a low but constant dose of CNO (∼0.22 mg/kg/h). Behavioral testing of male mice was initiated 12 days after minipump insertion and continuous CNO exposure using the test sequence depicted in Figure 1A and sFigure 1A. We used the EPM, OFT, and NSFT to assess changes in locomotion and negatively regulated, anxiety-related behavior under low and comparatively high acute stress conditions, respectively. In addition, we used the FUST to assess changes in positively regulated motivated behavior and hedonic drive in male mice. The sniffing of urine in this test is associated with dopamine release in the nucleus accumbens, indicating it is rewarding ^35^. Lastly, these initial experiments included the FST as a test traditionally used to assess antidepressant drug activity in rodents. The behavior of PLC-injected male mice in the EPM (Figure 1B) and OFT (sFigure. 1B) was unaffected by hM3Dq/CNO-mediated chronic activation of SST cells, indicating unaltered locomotion and anxiety-related behavior under low acute stress conditions. However, in the FST and NSFT, hM3Dq injected mice showed a reduced immobility score and a reduced latency to feed while in the FUST, they showed an increased time sniffing urine compared to empty vector controls, Thus, chronic activation of SST neurons in the PLC of male mice resulted in antidepressant drug-like changes in both positively and negatively regulated forms of motivated behavior.

**Figure 1.**
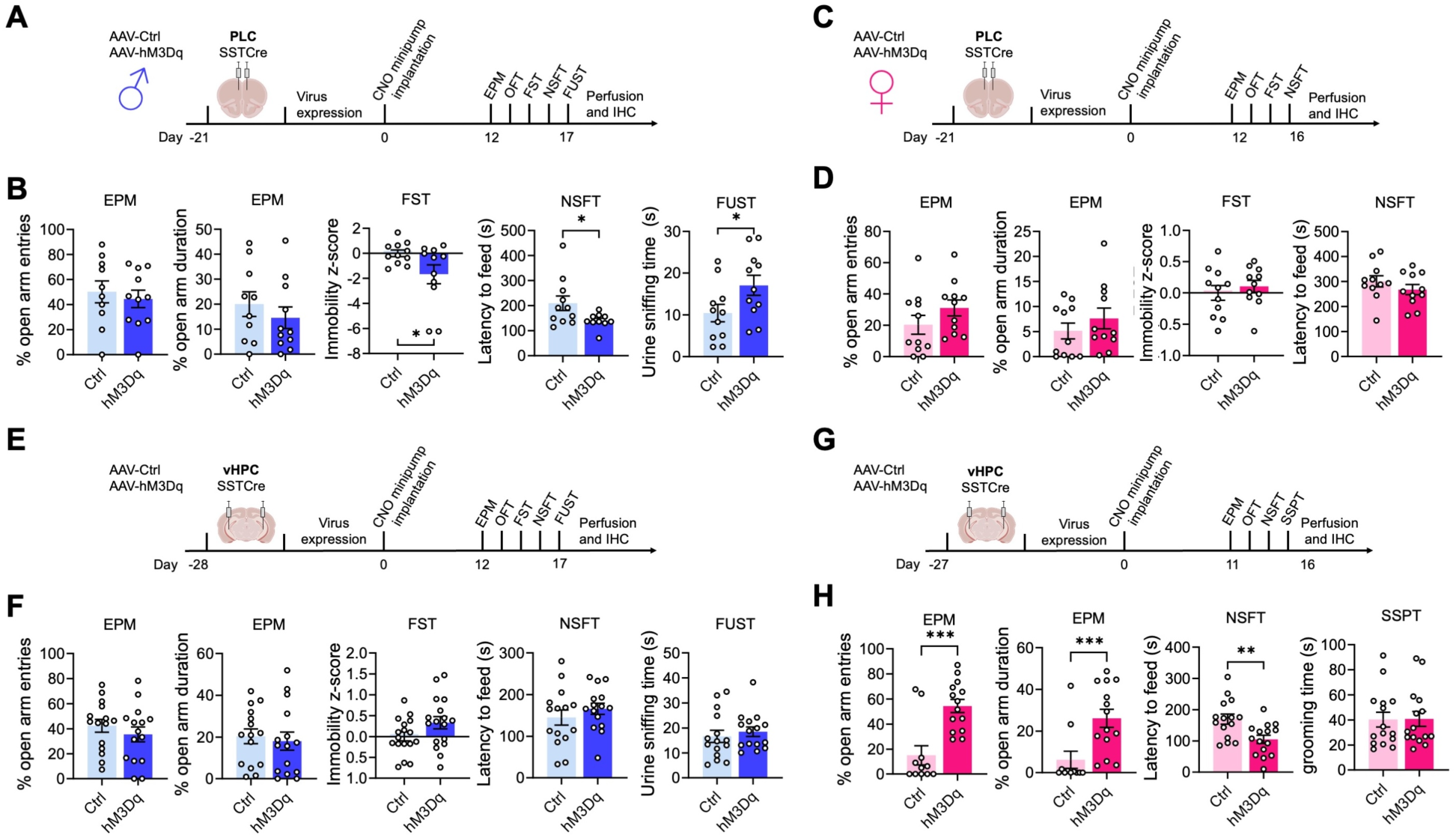
Chronic activation of SST neurons induces antidepressant-like changes in motivated behavior via sex-specific brain substrates. A–D) Chemogenetic manipulation of PLC-injected male (A, B) and female (C, D) mice including experimental designs (A, C) and results from the EPM, FST, and NSFT (both sexes) and FUST for males (B, D). Note that hM3Dq-mediated chronic activation of SST cells reduced the immobility z-score in the FST, the latency to feed in the NSFT, and increased the urine sniffing time in the FUST of male mice (p < 0.05, t-tests, for all comparisons) (B) but had no effect in the NSFT and FST of females (D). The behavior in the EPM was unaffected independent of sex (B, D). Results for the OFT are shown in sFigure 1. **E–H**) Analogous manipulation of male (E, F) and female (G, H) mice in the vHPC. hM3Dq-mediated activation of SST neurons in the vHPC had no effect on motivated behavior in males across all five tests (F) (t-tests) but increased the % open arm entries and % time on open arms (p < 0.001, Mann-Whitney) and reduced the latency to feed in the NSFT of female mice (p < 0.01, t-test). The SSPT substituted for the FST in this last experiment but was unaffected by SST neuron activation (see text). Means ± SEM. *p < 0.05, **p < 0.01, ***p < 0.001, n = 11–15, Mann Whitney, or t-test.

Next, we analogously tested the effects of PLC SST neuron activation in female mice (Figure 1C, D, sFigure 1C, D). The behavior in the EPM and OFT was unaffected as in males (Figure 1D, sFigure 1D). However, female mice differed from males in that activation of SST cells in the PLC had no effects in the NSFT and FST. This finding was unexpected because SSTCre:γ2^f/f^ mice with disinhibited SST neurons exhibit anxiolytic and antidepressant-like changes in behavior independent of sex ^36^. The data suggested that GABAergic control of motivated behavior in the PLC is male-specific and predicted that GABAergic control of motivated behavior in female mice is controlled by a distinct brain substrate.

### Chronic activation of SST neurons in the ventral hippocampus has anxiolytic and antidepressant-like effects on motivated behavior in female but not male mice

In search for a substrate for SST neuron-mediated changes in motivated behavior of female mice, we next focused on the ventral hippocampus (vHPC) as a second major brain region implicated in the top-down control of motivated behavior (Figure 1E–H, sFigure 1E–H). Interestingly, we found that the sex-specific effects of hM3Dq/CNO-mediated activation of SST neurons in the vHPC were opposite to those in the PLC, as indicated by the lack of an effect of SST neuron activation in the EPM, FST, NSFT, and FUST of male mice (Figure 1 E, F), and robust anxiolytic-like changes in the EPM and NSFT of female mice (Figure 1G, H). As a substitute for the FUST and FST, we subjected the vHPC-manipulated female mice to a Sucrose Splash Test. Grooming behavior in this test is accompanied by dopamine release in the nucleus accumbens ^34^, indicating that the behavior is rewarding similar to sniffing female urine for male mice. However, the behavior of hM3Dq/CNO manipulated female mice in the SSPT was unaffected in this experiment (sFigure 1H). In retrospect, this is consistent with recent evidence that dopamine release and grooming behavior in this test is selectively stimulated after chronically stressful experiences ^34^, as we confirmed further below. Notably, the anxiolytic-like effect of SST neuron activation in the vHPC of females was seen both under low-stress (EPM) and comparatively high-stress conditions (NSFT), whereas in PLC-manipulated male mice it was only seen under high-stress conditions (NSFT). Behavior in the OFT was unaffected, independent of sex and brain region manipulated (sFigure 1). Moreover, home cage feeding of food-deprived vHPC-manipulated animals was unaffected indicating that the behavioral changes in the NSFT were not due to an altered feeding drive. Collectively, these experiments indicated a striking sex-specific dissociation in the top-down control of antidepressant- and anxiolytic-like motivated behavior by prelimbic and hippocampal GABAergic microcircuits.

### SST neurons of the PLC and vHPC mediate resilience to chronic stress-induced defects in motivated behavior in male and female mice, respectively

We next asked whether activation of SST neurons in the PLC of male mice is sufficient to confer resilience to chronic stress-induced defects in motivated behavior. Virus-transduced mice were allowed to recover and subjected to an initial FUST to assess baseline behavior, followed by CNO minipump implantation, 15 days of CVS, minipump removal, and a battery of tests assessing negatively and positively motivated behavior including a second FUST (Figure 2A). The weight of mice during the three weeks of CVS was measured to assess chronic stress effects on whole-body physiology (Figure 2B). Interestingly, CVS resulted in a time-dependent body weight reduction in control but not hMD3q-injected mice, suggesting whole-body physiological stress protection. Repeated testing in the FUST before and after CVS revealed an overall stress effect independent of the vector injected, as well as an interaction between CVS and the type of vector injected. CNO reversed the CVS effect in hM3Dq-but not control vector-manipulated mice (Figure 2C). Analyses after CVS exposure further showed that hM3Dq-mediated activation of SST neurons had anxiolytic-like effects with regard to open arm entries and open arm duration in the EPM, and latency to feed in the NSFT. Collectively, these data indicated that SST neurons of the PLC were sufficient to mediate resilience to CVS-induced defects in both positively and negatively regulated behavior of male mice.

**Figure 2.**
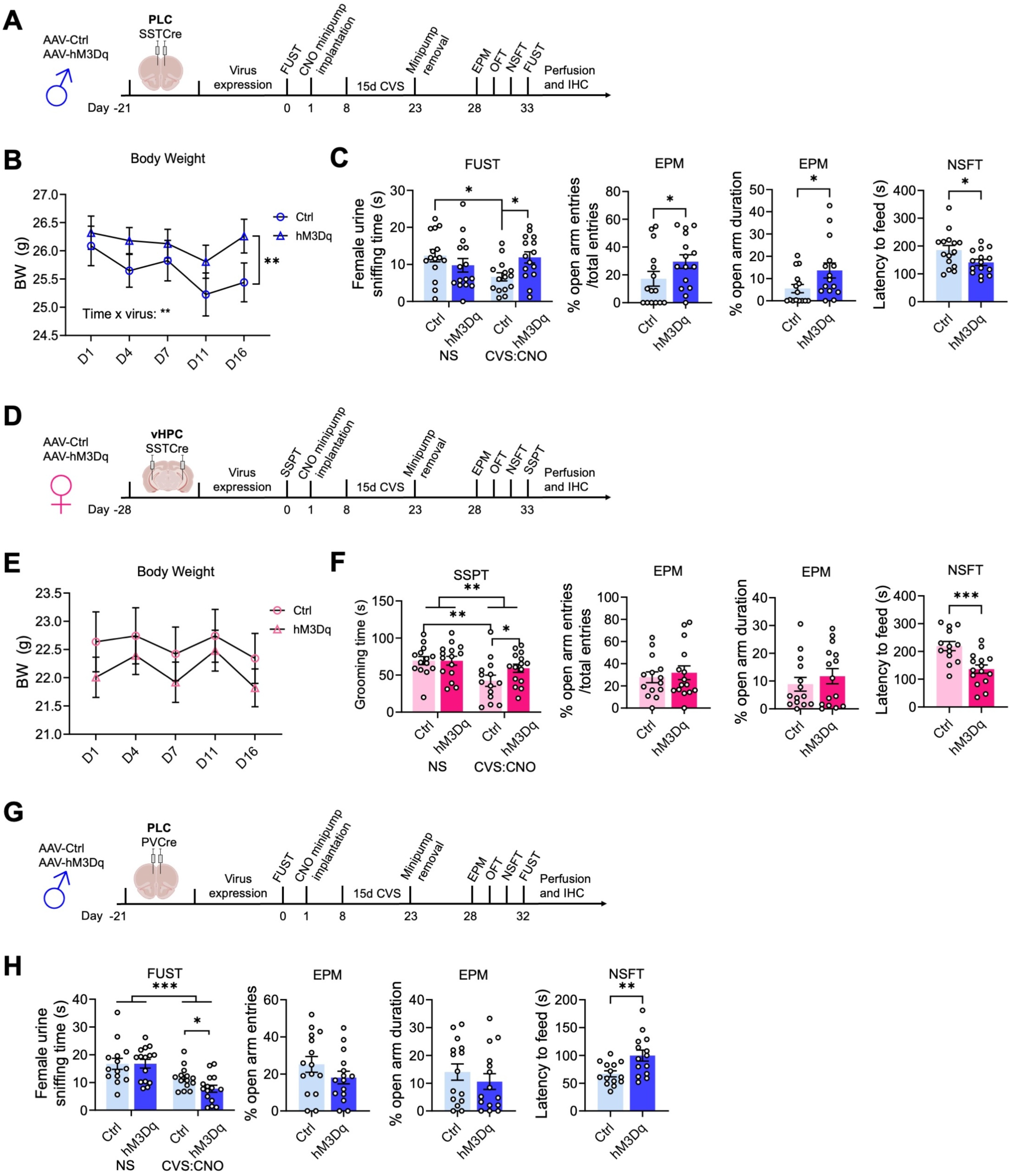
Activation of SST but not PV neurons induces resilience to chronic stress-induced changes in motivated behavior. A, D, G) Time course of manipulations of male (A, G) and female mice (D). The mice were injected with hM3Dq or control virus in the brain region indicated, followed by the analysis of baseline behavior in a FUST [males (A, G)] or SSPT [females (D)], implantation of CNO minipumps, exposure to 15 days of CVS, removal of minipumps, and testing in the EPM, OFT, NSFT, and a second FUST or SSPT. B) The weight of PLC-injected male mice showed a time x virus interaction (F_(4, 112)_ = 3.7, p < 0.01, RM ANOVA) with a time-dependent weight loss in control- but not hM3Dq-injected mice (p <0.01, Sidak test). **C**) Comparing the female urine sniffing times between the first and second FUST revealed a stress x vector interaction (F_(1,28)_ = 7.4, p < 0.05, RM ANOVA) that was significant for control- but not hM3Dq vector-injected mice (p < 0.05, Sidak test). hM3Dq vs. control injected, CVS-exposed mice showed a significantly greater urine sniffing time (p < 0.01). In addition, hM3Dq- vs. control-injected CNO- and CVS-exposed mice showed anxiolytic-like changes in motivated behavior in the EPM and NSFT (p < 0.05, t-tests, for all comparisons). **E**) vHPC-injected female mice exposed to CVS showed no measurable change in body weight independent of vector injected. **F**) The time spent grooming in the SSPT showed an overall stress effect (F_(1, 27)_ = 9.799, p < 0.01, RM ANOVA) with a significant reduction in control- (p < 0.01) but not hM3Dq-injected mice. A planned comparison of the grooming times after CVS showed an increase in the grooming duration in hM3Dq- vs. control injected mice (p < 0.05). In addition, in the NSFT hM3Dq-injected mice showed lower latency to feed compared to control-injected mice (p < 0.001, t-test). **G**) Analyses of PVCre male mice injected in the PLC analogous to SSTCre males in (B) and (C). **H**) In the FUST, a comparison of the urine sniffing times before and after CVS exposure showed a stress effect independent of the vector injected (F_(1, 27)_ = 33.96, p < 0.001, RM ANOVA). A planned comparison of control- and hM3Dq-virus-injected mice after CVS exposure revealed a reduction in the urine sniffing time of hM3Dq- vs. control-injected mice (p < 0.05). The % open arm entries and open arm times in the EPM of CVS-exposed mice remained unaffected by virus treatment. However, in the NSFT, hM3Dq- vs. control-injected mice showed an anxiogenic-like increase in the latency to feed (p < 0.01). Data represent means ± SEM, *p < 0.05, **p < 0.01, ***p < 0.001, 2-way ANOVA plus Sidak test or t-test.

Next, we similarly tested whether SST neurons of the vHPC confer stress resilience to female mice (Figure 2D). In contrast, to males, CVS failed to affect the weight of female mice independent of DREADD manipulation (Figure 2E). Despite this qualitative sex difference in the stress effect, comparing the behavior in the SSPT before and after CVS revealed a significant CVS effect independent of virus (Figure 2F, first panel). Post hoc analyses showed that this effect was significant in control vector- but not hM3Dq vector-injected mice. As in PLC-manipulated males, hM3Dq-injected female mice showed an anxiolytic effect in the NSFT (Figure 2F, fourth panel). Therefore, SST neurons of the vHPC mediate resilience to CVS-induced defects in positively and negatively regulated behavior in female mice. Notably, SST neuron activation in the vHPC of females had no effect on % open arm times and % open arm entries in the EPM (Figure 2F, second and third panel), in contrast to SST neuron activation in the PLC of males (Figure 2C), and the behavior in the OFT and home cage feeding of food-deprived mice was unaffected, independent of sex, brain region and interneuron type (sFigure 2A–F). These experiments suggest that the brain regions that mediate sex-specific antidepressant-like behavioral effects in the absence of stress also confer resilience to the detrimental behavioral consequences of chronic stress exposure. In male mice, GABAergic PLC-mediated stress resilience extends to stress-induced weight loss, whereas in female mice this parameter could not be addressed due to the absence of a significant stress effect.

### Chronic activation of parvalbumin-positive interneurons in the PLC has detrimental effects in male mice

Unlike the pro-resilience function of SST neurons in the PLC of male mice, increased activity of parvalbumin (PV)-positive interneurons may have detrimental effects, as shown by others for the mPFC of female mice and hippocampus of male mice ^37, 38^. To assess the function of PV neurons in the PLC of male mice, we here assessed the behavioral consequences of chemogenetic PV neuron activation in CVS-exposed mice. Comparing the female urine sniffing times before and after CVS revealed a significant stress effect independent of virus injected, as expected. However, in contrast to the pro-resilience effect of PLC SST neuron activation (Figure 2C), activation of PLC PV neurons exacerbated the CVS effect and led to a reduction in sniffing time in hM3Dq- vs. control vector-injected, CVS-exposed mice (Figure 2H). Moreover, in contrast to the anxiolytic-like effect of PLC SST neuron activation in the NSFT (Figure 2C), activation of PLC PV neurons during CVS exposure had anxiogenic-like effects, as evidenced by the additional increase in the latency to feed in hM3Dq- vs. control virus-injected mice (Figure 2H). Thus, the pro-resilience effect of GABAergic inhibition is specific to dendrite targeting GABAergic inhibition and not observed with PV-neuron mediated inhibition that targets the somatodendritic compartment of pyramidal cells. Notably, the behaviors of hM3Dq-transduced PVCre mice in the EPM (Figure 2H) and OFT (sFigure 2F) were unaltered, suggesting that the anxiogenic-like effects of PV neuron activation are manifested selectively under acutely stressful conditions.

### SST neurons in the PLC and vHPC act sex specifically to reverse chronic stress-induced defects in motivated behavior in male and female mice, respectively

We next asked whether activation of SST neurons in the PLC would reverse prior CVS-induced defects in motivated behavior. Male mice transduced with hM3Dq or control virus were subjected to an initial FUST to assess baseline behavior, followed by three weeks of CVS and a second FUST to verify the stress effects (Figure 3A). We then implanted CNO minipumps, followed 12 days later with a final behavioral test battery, including a third FUST. To prevent the possible fading of stress effects during the extended time course of experimentation we preceded the final behavioral assessment by three days of CVS exposure, which by itself does not affect motivated behavior ^30^. Comparison of PLC control- and hM3Dq-vector-injected mice in the FUST before CNO treatment revealed an overall stress effect, independent of the vector injected (Figure 3B, 1^st^ panel). Comparison of CVS-exposed mice before and after CNO treatment in the second and third FUST revealed an overall CNO effect with a significantly greater Female Urine Sniffing Time in hM3Dq vs. control injected mice (Figure 3B, 2nd panel). Thus, chronic activation of PLC SST neurons effectively reversed the anhedonia-like consequences of CVS exposure. The behavior in the EPM and OFT remained unaffected, indicating unaltered locomotion and anxiety-related behavior under low-stress conditions (sFigure 3A, B). However, the latency to feed in the NSFT (a comparatively high-stress situation) was significantly reduced in hM3Dq- vs. control-vector-injected mice, indicating a reversal of stress-induced anxiety-like behavior. The home cage feeding of hM3Dq-injected mice was reduced, indicating that the change in the latency to feed was not explained by an altered feeding drive (sFigure 3B, last panel).

**Figure 3.**
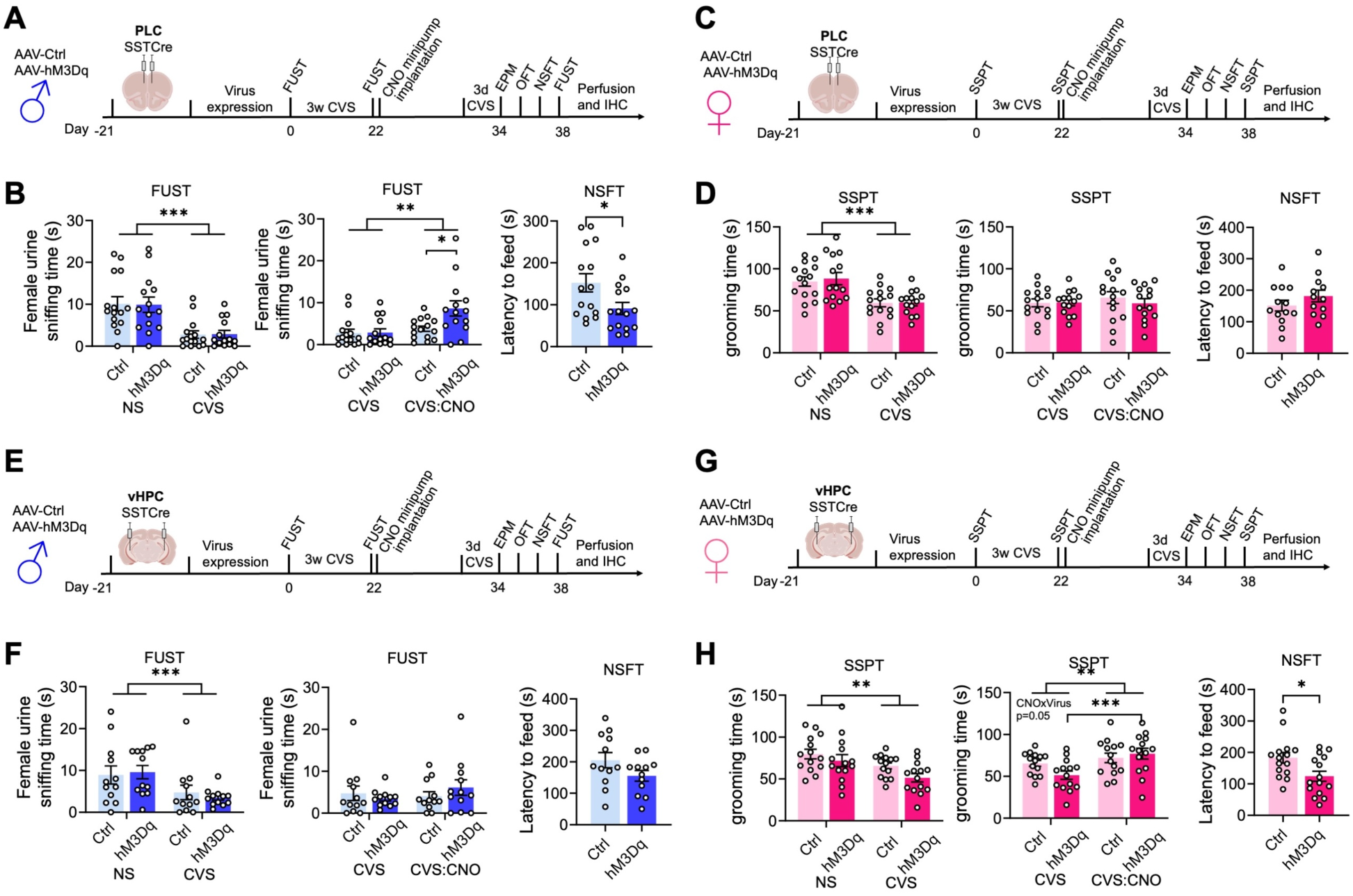
Activation of somatostatin neurons reverses the behavioral effects of chronic stress exposure through sex-specific brain substrates. A, C, E, G) Time course of manipulations of male (A, E) and female mice (C, G). The mice were injected with hM3Dq or control virus in the brain region indicated, followed by analysis of baseline behavior in a FUST [males (A, E) or SSPT (females (C, G)], exposure to 21 days of CVS, a second FUST or SSPT respectively, implantation of CNO minipumps, 3 days of CVS, and testing in the EPM, OFT, NSFT, and a third FUST or SSPT, respectively. **B**) Comparing the female urine sniffing times of PLC-injected male mice in sequential FUSTs before and after CVS revealed an overall stress effect (F_(1,27)_ = 35.96, p < 0.001, RM ANOVA) independent of vector injected. Comparing the female urine sniffing times of CVS-exposed mice before and after CNO revealed an overall CNO effect (F_(1,27)_ = 9.399, p < 0.01) and a vector x CNO interaction (F_(1,27)_ = 4.640, p < 0.05, RM ANOVA). CNO increased the urine sniffing time in hM3Dq- vs. control-injected mice (p < 0.05). The latency to feed of hM3Dq vs. control injected was reduced (p < 0.05). **D**) Comparing the grooming times of PLC-injected female mice in sequential SSPTs before and after CVS revealed an overall stress effect (F_(1,27)_ = 25.07, p < 0.001, RM ANOVA). CNO did not affect the grooming time independent of the vector. The latency to feed in the NSFT after CVS and CNO was unaffected. **F**) Comparing the female urine sniffing times of vHPC-injected male mice before and after CVS revealed an overall stress effect (F_(1,27)_ = 19.55, p < 0.001, RM ANOVA) independent of the vector injected. CNO did not affect the grooming time independent of the vector. The latency to feed was unaffected by the vector injected. **H**) Comparing the grooming times of vHPC-injected female mice in sequential SSPTs before and after CVS revealed an overall stress effect (F_(1,26)_ = 11.74, p < 0.01, RM ANOVA). Comparing the grooming times of the CVS-exposed mice before and after CNO revealed an overall CNO effect and a CNO x vector interaction (F_(1,26)_ = 4.064, p = 0.05, RM ANOVA). CNO increased the grooming time of CVS-exposed hM3Dq- (p < 0.001) but not control-injected mice. The latency to feed after CVS and CNO exposure of hM3Dq- vs. control-injected was reduced. Data represent means ± SEM, *p < 0.05, **p < 0.01, ***p < 0.001, 2-way ANOVA, Sidak, or t-test.

Analogous testing of female mice and substitution of SSPTs for the FUSTs and comparing the grooming time before and after CVS exposure revealed a significant stress effect independent of the vector injected (Figure 3C, D, first panel). However, CNO did not affect the grooming time, independent of the vector injected (Figure 3D, 2^nd^ panel). Similarly, a pairwise comparison of control and hM3Dq injected mice after CVS and CNO exposure in the NSFT showed no difference between control and hM3Dq injected mice. Therefore, in contrast to male mice, activation of SST neurons in the PLC of female mice failed to reverse CVS-induced anhedonia- and anxiety-like behavior. Notably, the behavior in the OFT and EPM was unaltered and consistent with unaltered locomotion (sFigure 3C, D).

The findings thus far suggested that similar to effects on motivated behavior in the absence of stress, the reversal of chronic stress effects by GABAergic inhibition was sex and brain region-specific. To further establish this, we tested the ability of GABAergic circuits in the vHPC to reverse the effects of prior stress exposure (Figure 3 E, F). We injected the vHPC of SSTCre mice with control or hM3Dq virus, exposed the mice to CVS, and then implanted CNO-releasing minipumps (Figure 3E). Comparison of male mice in the FUST before and after CVS exposure revealed an overall stress effect, independent of virus transduced, as expected (Figure 3F, first panel). However, in contrast to manipulation of the PLC, CNO exposure of hmD3q injected animals failed to reverse the CVS effect (Figure 3F, 2^nd^ panel). Similarly, a comparison of control- and hM3Dq-injected mice after CVS and CNO exposure failed to show anxiolysis in the NSFT of hM3Dq-injected mice (Figure 3F, 3rd panel). By contrast, analogous testing of female mice injected in the vHPC (Figure 3G) and analyzed in the SSPT revealed a significant CVS effect that was reversed by CNO exposure selectively in hM3Dq- but not control-injected mice (Figure 3H, 1^st^ and 2^nd^ panel). Moreover, testing in the NSFT revealed an anxiolytic-like effect of CNO selectively in hM3Dq but not control-injected mice (Figure 3H, 3rd panel). Thus, the reversal of chronic stress-induced anxiety and anhedonia-like behavior in male and female mice mediated by GABAergic microcircuits involves distinct sex-specific cortical substrates.

### SST neuron-mediated reversal of CVS effects on motivated behavior involves increased activity of pyramidal cells

GABAergic synaptic transmission is widely known to inhibit glutamatergic depolarizing inputs and hence to reduce dendritic spiking and output of glutamatergic target neurons ^39^. By contrast, all evidence suggests that antidepressant drug-like changes in motivated behavior involve increased rather than reduced activity of cortical output neurons ^18, 20, 21^. We hypothesized, therefore, that chronically and moderately elevated GABAergic transmission at dendrites may lead to increased rather than reduced activity of pyramidal target cells. To test this prediction, we quantitated the densities c-Fos and FosB-expressing neurons as proxies for cell-specific neural activity in vivo. Male mice that had been transduced with AAV-hM3Dq or control virus in the PLC, followed by CVS exposure and CNO-induced reversal of behavioral stress effects (Figure 3A, 4A), were subjected to immunostaining for c-Fos and virus-encoded, Cre-dependent mCherry to assess the density of c-Fos^+^, mCherry^+^ cells representing SST cells, as well c-Fos^+^ non-SST (mCherry-negative) cells representative of pyramidal cells in areas exhibiting virally transduced SST cells. Precisely 90 min before euthanasia the mice were exposed to a novel arena for 10 min to induce a mild but behaviorally relevant arousal state (Figure 4A). The density of c-Fos^+^ SST neurons was increased in hM3Dq- compared to control virus-transduced mice, which confirms that chronic low-level CNO-delivery increased the activity of hM3Dq expressing SST neurons (Figure 4 B1,2) with 18.7% and 29.0% of mCherry+ SST neurons showing c-Fos expression in ctrl and hM3Dq injected mice, respectively. Importantly, the density of c-Fos^+^ non-SST (mCherry-negative) cells was increased in parallel (Figure 4 B3), indicating that chronically increased inhibition by SST cells leads to paradoxically increased activation of non-SST cells. The density of c-Fos^+^ PV^+^ cells was less than 5% of the density of c-Fos^+^ non-SST cells, indicating that c-Fos^+^ non-SST cells consist predominantly of pyramidal cells. To corroborate these findings, we quantitated the density of mCherry+ SST neurons expressing FosB, a marker that is induced by neuronal depolarization and Ca^2+^ entry similar to c-Fos but with a slower and longer-lasting time course ^40^. The density of FosB^+^ SST cells was increased in hM3Dq vs. control injected mice, with 7.9% and 20.2% of SST neurons showing expression of FosB in control and hM3Dq virus injected mice, respectively (Figure 4C). Moreover, the density of FosB^+^ non-SST cells was increased in hM3Dq vs. control-injected mice (Figure 4C), thereby confirming that SST neurons increase the activity of their putative pyramidal target cells.

**Figure 4.**
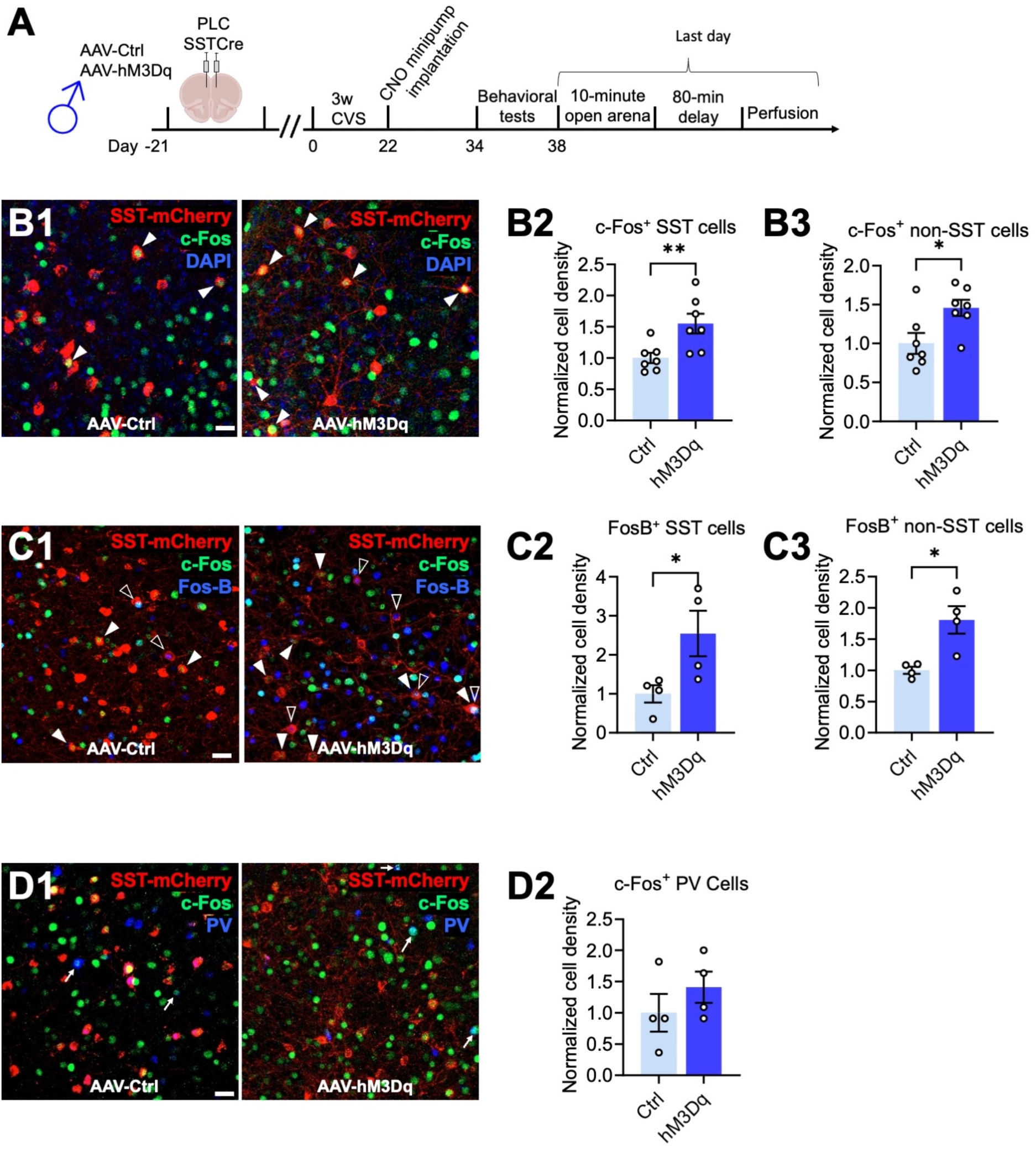
hM3Dq-mediated activation of SST cells in the PLC increases the density of c-Fos and FosB immuno-positive non-pyramidal cells. **A**) Time course of manipulations. On the day after the last day of behavioral testing, the mice were exposed to a novel open arena for ten minutes and the brain perfused 80 minutes later. **B**) Triple staining for virus-expressing SST neurons (mCherry, red), c-Fos (green) and the nuclear stain, DAPI (blue) with representative micrographs of control- (left) and hM3Dq- (right) infected animals (B1) and summary data showing increased densities of c-Fos^+^ SST (B2, p < 0.01) and c-Fos^+^ non-SST neurons (p < 0.01, n = 7) in hM3Dq- vs. control-injected mice. **C)** Triple staining for SST neurons for mCherry (red), c-Fos (green), and FosB (blue) with representative micrographs of a control- (left) and hM3Dq-injected animals (right) (C1) and summary data showing increased densities of FosB^+^ SST (C2, p < 0.05) and c-Fos^+^ non-SST neurons (p < 0.05, n = 4) in hM3Dq- vs. control-injected mice. **D**) Triple staining for SST neurons (mCherry, red), c-Fos (green), and PV (blue) with representative micrographs of control- (left) and hM3Dq-infected animals (right) (D1) and summary data showing unaltered densities of c-Fos^+^ PV cells (n = 4). The densities of c-Fos^+^ and FosB^+^ SST cells were normalized to the total densities of virally transduced SST cells. Solid arrowheads (B1) point to c-Fos^+^,mCherry^+^ cells; empty arrowheads (C1) point to FosB^+^,mCherry^+^ cells, and arrows (D1) point to c-Fos plus FosB double positive cells. Scale bar, 30 μm. Data represent means ± SEM, *p < 0.05, **p < 0.01, t-test.

Optogenetic stimulation of the large majority of cortical SST neurons results in inhibition of pyramidal cells ^41^. However, a small subset of SST^+^ neurons in cortical layer IV has been shown to predominantly inhibit PV^+^ cells ^41^, which raised the question of whether the SST neuron-mediated activation of pyramidal cells reported here involved disinhibition via PV cells. We found that the density of c-Fos and PV double-positive neurons was unaltered (Figure 4D), indicating that SST neuron-mediated activation of pyramidal cells cannot be explained by PV neuron-mediated disinhibition. It follows, that the antidepressant-like, SST neuron-mediated reversal of chronic stress-induced changes in motivated behavior can only be explained by direct SST neuron-mediated activation of pyramidal cells. A proposed mechanism is discussed.

## DISCUSSION

We here have used a novel chronic chemogenetic approach to map GABAergic neural circuits that mediate antidepressant drug-like behavioral changes, including resilience to the anxiogenic and anhedonia-like changes of chronic stress and reversal of chronic stress-induced behavioral effects. As a first major finding, we report that SST neurons in the PLC mediate such responses in males but not females, while functionally homologous SST interneurons in the vHPC mediate such responses in female but not male mice. In contrast to the pro-resilience effects of SST neurons, we found that increased PV neuron activity has detrimental effects in that it exacerbates anxiety and anhedonia-like behavior in stress-exposed animals. Long-range inputs from mPFC ^42^ and vHPC ^43^ to the amygdala and nucleus accumbens are known to contribute to the top-down control of anxiety and anhedonia-related changes in motivated behaviors, and they are both known to be highly sensitive to chronic stress exposure. The mPFC and vHPC are also similarly vulnerable to stress and known as key substrates mediating the detrimental effects of chronic stress exposure. Our data reveal striking sex differences in the relative importance of GABAergic microcircuits in the mPFC and vHPC and/or their subcortical target areas. Such sex differences are likely to be important for qualitative differences in the sex-specific vulnerabilities and resilience to stress-induced anxiety- and anhedonia-like motivated behavior.

As a second major finding, we report that the antidepressant-like, CVS-reversing behavioral consequences of SST neuron activation are associated with increased rather than reduced activity of their putative pyramidal cell targets. This was surprising because there is plenty of evidence that SST neurons inhibit their target cells. First, the antidepressant-like behavioral phenotype of mice with disinhibited SST neurons is associated with strengthened inhibitory inputs onto pyramidal cells ^24^. Second, acute stimulation of Martinotti-type SST cells is widely known to hyperpolarize pyramidal cells ^41^ and to reduce dendritic Ca^2+^ influx ^39^ ^44, 45^, while acute inactivation of Martinotti cells leads to disinhibition and action potential firing of pyramidal cells ^41^. However, in support of our findings, optogenetic stimulation of glutamatergic neurons in the mPFC is known to mimic antidepressant drug-induced behavioral effects ^18–21^. Moreover, a tonic form of GABAergic inhibition at pyramidal cell dendrites mediated by α5-GABA_A_ receptors has been shown to de-inactivate low threshold V-gated Ca^2+^ channels and potentiate glutamatergic activity-induced Ca^2+^-influx and dendritic spiking of pyramidal cells ^46^. Lastly, α5-GABA_A_ receptors represent a promising target for next-generation antidepressants ^47, 48^. Therefore, antidepressant drug-like behavioral effects of *low-level* and *chronically* enhanced GABAergic inhibition at dendrites involve disinhibition of pyramidal cells. These data provide a mechanistic rationale for the lack of antidepressant efficacy of benzodiazepines, which almost exclusively potentiate GABA_A_ receptors underlying phasic inhibition, but indiscriminately for virtually all neurons in the CNS. In particular, we showed here that potentiation of PV neuron-mediated inhibition in the PLC has detrimental, pro-depressant-like consequences even in already compromised, chronically stressed animals.

The large majority of neocortical SST interneurons (layers 2/3 and 5) target the distal apical dendrites of pyramidal cells. However, a morphologically distinct subgroup of SST cells in cortical layer IV predominantly inhibits PV^+^ interneurons, which leads to disinhibition of principal cells ^41^ and raises the possibility that SST-neuron-mediated activation of pyramidal cells involves PV-neuron mediated disinhibition. However, when we used c-Fos as a proxy for neural activity in behaving animals we found no evidence for SST neuron-mediated inhibition of PV neurons. Moreover, stress-induced changes in the activity of PV cells in the mPFC have been shown to alter the emotional reactivity of female but not male mice ^37^ which further indicates that PV cells are unlikely to contribute to the mPFC-dependent changes in behavior of male mice in our study. Of note, when designing our chemogenetic stimulation protocol we reasoned that drastically increasing GABAergic dendritic inhibition would interfere with plasticity at dendrites needed to increase the activity of target cells and, accordingly, chose a low concentration of CNO (∼0.2 mg/kg/h) that is at, if not below, the threshold required for measurable hM3Dq-induced changes in neural activity in vitro ^49^. However, future experiments will need to establish the optimal level of SST neuron activation for most effective reversal of chronic stress effects.

## CONCLUSION

Benzodiazepines which inhibit the activity of neurons indiscriminately along the neural axis and preferentially at GABAergic synapses (phasic inhibition), are widely known to be ineffective as antidepressants ^50^. By contrast, selective enhancement of tonic GABAergic inhibition at dendrites through α5-GABA_A_ receptors ^51^ (perhaps with contribution of α2-GABA_A_ receptors ^52^) holds significant promise as target for antidepressant drug development ^47^. We propose that SST neurons are acutely neuro-protective during stress exposure and very high network load by limiting NMDA receptor-mediated Ca^2+^ influx into dendrites via activation of prototypical postsynaptic GABA_A_ receptors ^44^ and by facilitating recovery of pyramidal cells in between stressful periods through potentiation of tonic inhibition, which increases excitability of pyramidal cells. Indeed, tonic inhibition through α5-GABA_A_ receptors at dendrites leads to de-inactivation of t-type Ca^2+^ channels and thereby increases dendritic excitability and pyramidal cell output ^46^. By contrast, acute potentiation of GABAergic synaptic inputs to dendrites reduces Ca^2+^ influx to dendrites and thus exerts an inhibitory function on pyramidal cells ^44^. Importantly, while the brain substrate for GABA-mediated neuroprotective and antidepressant effects is strikingly sex-specific, similar properties of SST neurons in the PLC and vHPC, along with the shared subcortical targets of these brain regions, should allow for α5-GABA_A_ receptor-mediated antidepressant therapies to be efficacious independent of sex.

## Acknowledgments

We thank Yao Guo for her expert technical assistance. This publication was made possible by a grant (MH099851) from the National Institute of Mental Health (NIMH) to B.L. and generous support from Penn State University. Its contents are solely the responsibility of the authors and do not necessarily represent the views of Penn State or of the NIMH.

