## Supplemental Data for "Sex-specific GABAergic microcircuits that switch vulnerability into resilience to stress and reverse the effects of chronic stress exposure"

Supplementary Figure 1

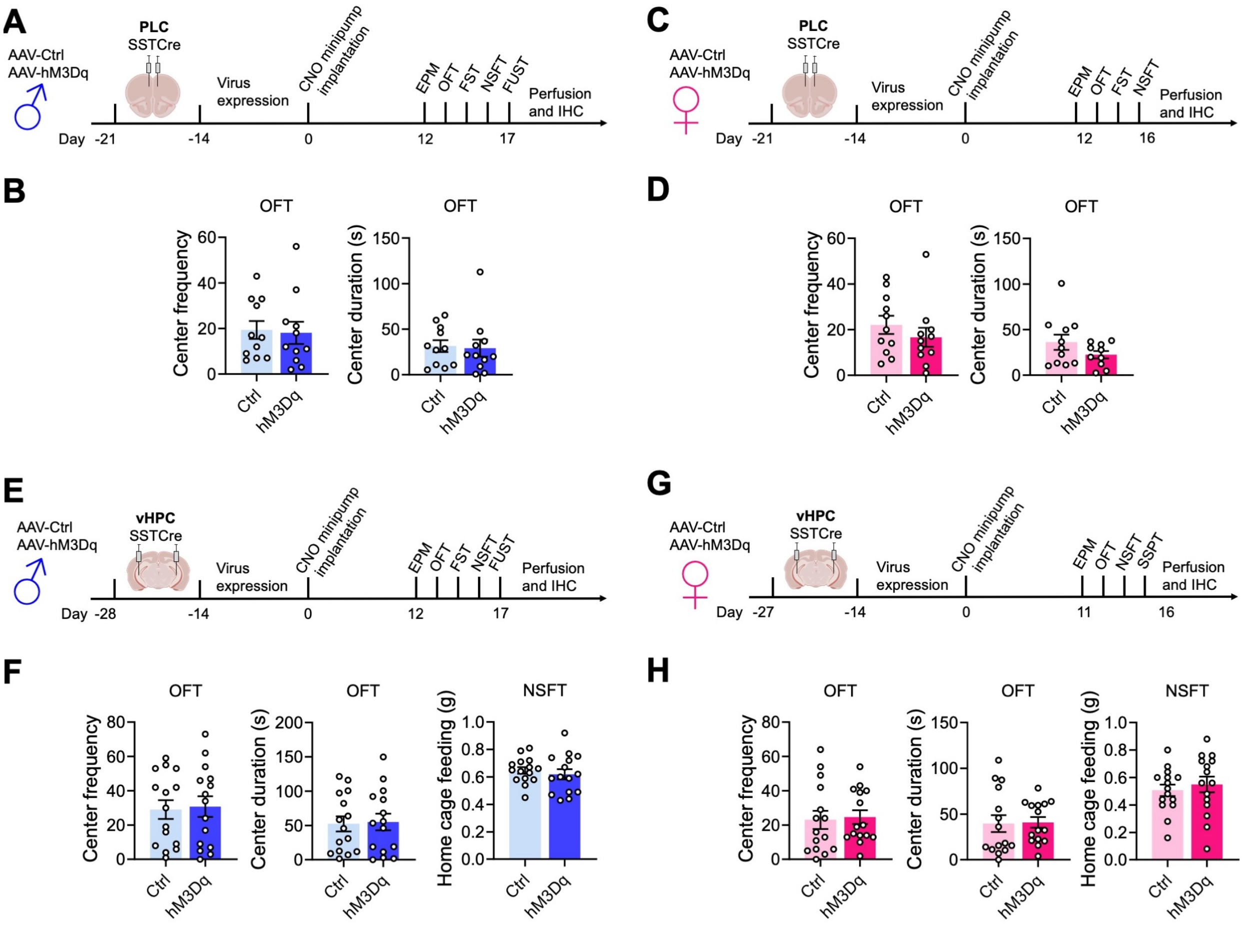


**sFigure 1. SST neuron-mediated changes in motivated behavior do not involve changes in locomotion or feeding drive.** A, C, E, G) Time course of experimentation and brain areas manipulated. B, D, F, H) Behavior in the OFT was unaffected by hM3Dq mediated activation of SST cells, independent of sex and brain region manipulated (B, D, F, and H). Home cage feeding of food-deprived mice was unaffected by SST neuron activation, as examined for vHPC-manipulated mice (F, H). Bar graphs represent means ± SE.

Supplementary Figure 2

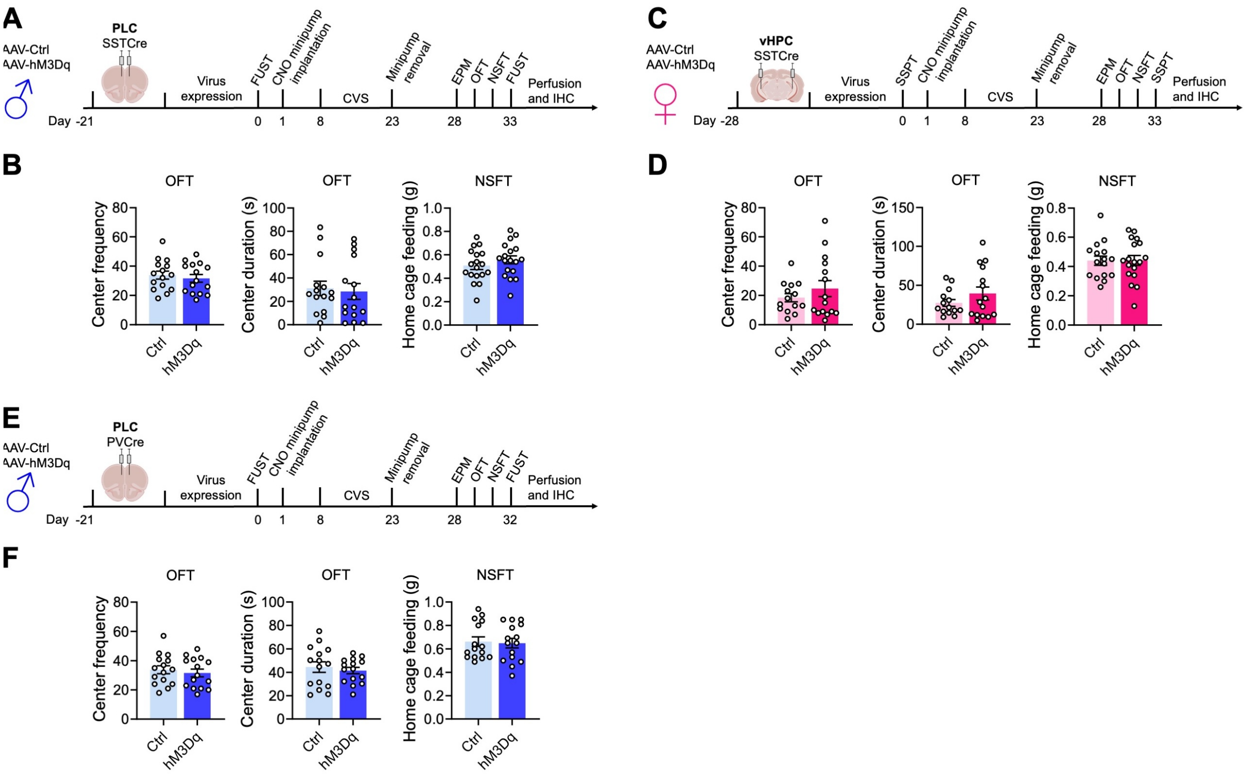


**sFigure 2. Activation of SST but not PV neurons induces resilience to chronic stress-induced changes in motivated behavior. A, C, G)** Time courses of manipulations of male (A) and female SSTCre mice (D) in the PLC and vHPC, respectively, and of PVCre mice on the PLC (E). The Behavior of hM3Dq vs. control virus-injected mice in the OFT and NSFT was unaltered in all three conditions. Data represent means ± SEM, t-test.

Suppl. Figure 3

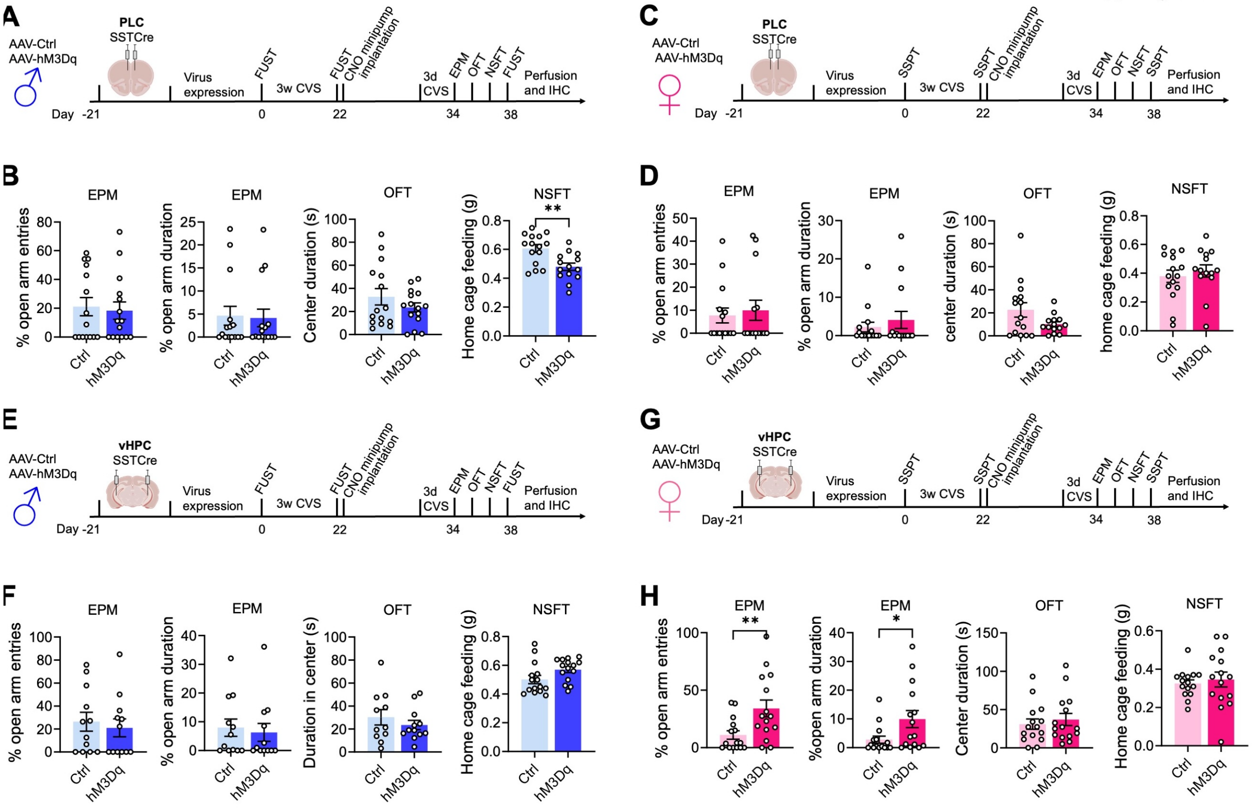


**sFigure 3. Activation of somatostatin neurons reverses the behavioral effects of chronic stress exposure through sex-specific brain substrates. A, C, E, G)** Time courses of manipulations of male (A, E)  and female SSTCre mice (D, G) in the PLC (A, C) and vHPC (E, G). The behavior of hM3Dq vs. control virus injected mice in the EPM (% open arm entries and time), OFT (center duration), and NSFT (home cage feeding) was unaltered in all experiments (B, D, E, G, F, H) except for the reduced home cage feeding in hM3Dq vs. control virus-injected mice in PLC injected male mice (B) which does not explain the reduced latency to feed depicted in the corresponding main figure. Data represent means ± SEM, t-test.
